# Regimes and mechanisms of transient amplification in abstract and biological networks

**DOI:** 10.1101/2021.04.01.437964

**Authors:** Georgia Christodoulou, Tim P. Vogels, Everton J. Agnes

## Abstract

We use upper triangular matrices as abstract representations of neuronal networks and directly manipulate their eigenspectra and non-normality to explore different regimes of transient amplification. Counter–intuitively, manipulating the imaginary distribution can lead to highly amplifying regimes. This is noteworthy, because biological networks are constrained by Dale’s law and the non-existence of neuronal self-loops, limiting the range of manipulations in the real dimension. Within these constraints we can further manipulate transient amplification by controlling global inhibition.

---

Recurrent network models are known to produce different types of dynamics, ranging from regular to irregular, and from transient to persistent activity^1–6^. Moulding network dynamics to resemble experimental observations usually involves changes in the network architecture, i.e., the existence of synapses and their efficacies^7–9^. With this approach, the eigen-spectrum and the non-normality of the connectivity matrix are indirectly affected, and the relationship between changes in those qualities of the weight matrix and the network dynamics remain nebulous. Here, we manipulate the spectrum and non-normality of upper triangular matrices, such that their characteristics can be directly translated into dynamical properties (Fig. 1A). These matrices no longer represent the neuronal connectivity, but modes of activation that are arranged in a feedforward manner^10–12^. We are particularly interested in the different forms of transient amplification, a phenomenon that can resemble motor cortex activity during reaching^13–15^ and also emulate long-lasting working memory dynamics^16–18^. After a dissection of the underlying mechanisms of transient amplification using general upper triangular matrices, we consider biological constraints on the spectral distributions, and consequently, on the dynamics. Finally, we show how we can implement our findings in a biological plausible connectivity matrix with excitatory and inhibitory neurons, i.e., a matrix satisfying Dale’s law.

**FIG. 1.**
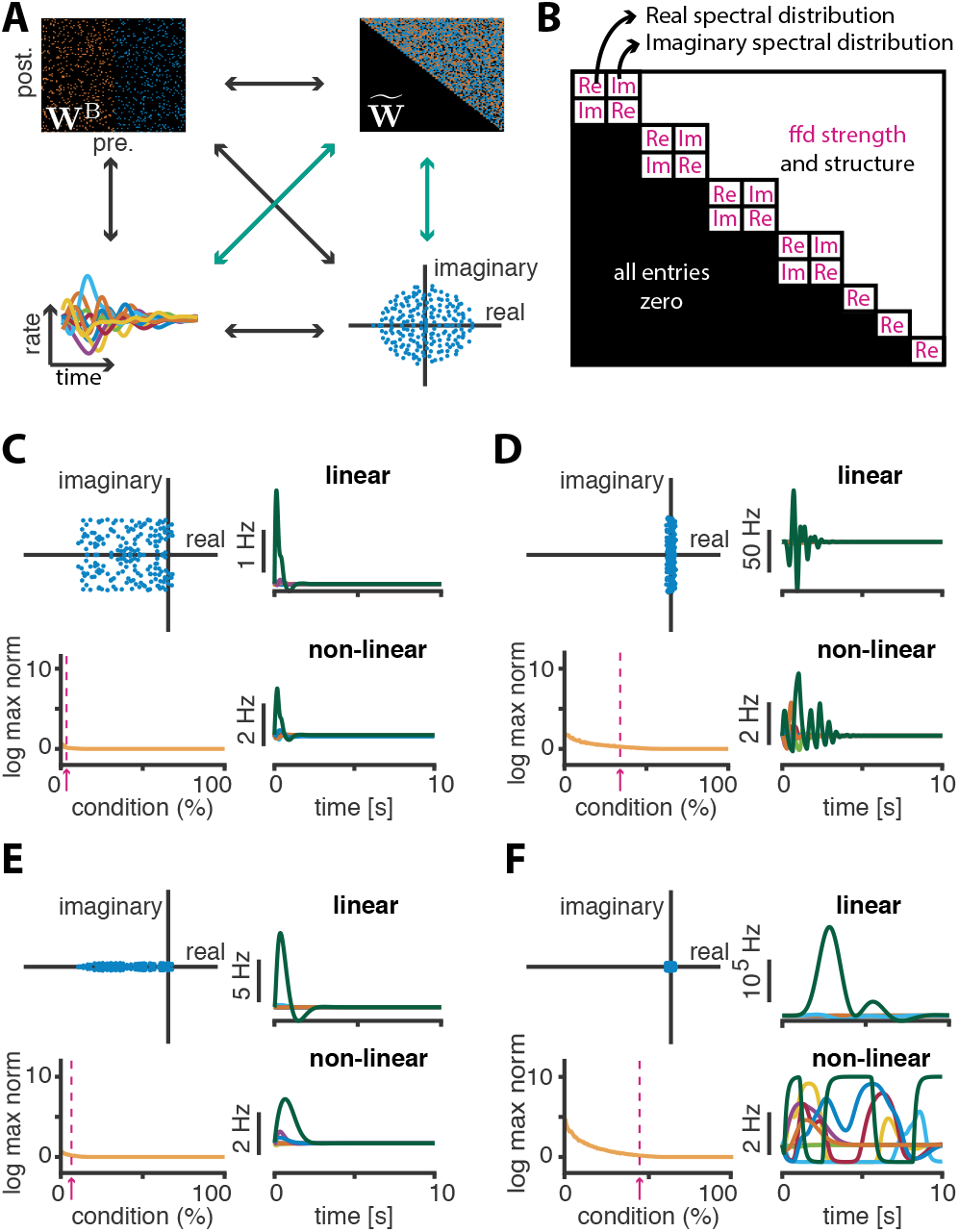
Eigenspectra and their respective dynamics. **A**, Schematic of the elements explored in this letter. Top left and clockwise: The connectivity matrix **W**^B^; its corresponding Schur upper triangular decomposition 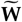; the eigenspectrum; and the induced dynamics. **B**, The upper triangular matrix 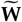 with the quantities that we alter in this letter in pink. **C–F**, Four cases of eigenspectra and their dynamics. In each panel clockwise: The spectrum; linear dynamics; non-linear dynamics; the logarithm of the maximum norm of the firing rate per initial condition. Pink dotted line and arrow correspond to the last initial condition whose norm is amplified by at least 50%. The feedforward structure is taken from a SOC^8^ and its Frobenius norm is fixed to 75. Real and imaginary parts follow an uniform distribution with diameters *d*_im_ and *d*_re_, respectively. **C**, When *d*_im_ = *d*_re_ = 10 only two conditions are slightly amplified. **D**, When *d*_im_ = 10 and *d*_re_ = 1, the system is capable of more amplification. **E**, Here *d*_im_ = 1 and *d*_re_ = 10 and surprisingly this also creates more amplification compared to the case shown in C. **F**, When *d*_im_ = *d*_re_ = 1, the system amplifies almost half of the initial conditions. The dynamics, given an initial condition of norm 1 reach the value of ~ 10^5^ in the linear case and consequently long-lasting dynamics in the non-linear case.

Throughout the paper we use the following notation for the connectivity matrix: **W** for a generic connectivity matrix, 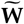 for a matrix given in upper triangular form, and **W**^B^ for a matrix following biological constraints. The dynamics of the recurrent network are defined by

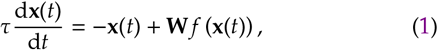

where **x**(*t*) is the internal state of the network at time *t*, and can be understood here as the membrane potential of a given neuron. This internal state of the neurons evolves with a characteristic time constant *τ* and is affected by the activity of other neurons of the network through the recurrent connections determined by **W**. Finally, the activation function, *f*(**x**(*t*)) = **r**(*t*), represents the input-output relation between the internal state, **x**(*t*), and the firing rate deviation, **r**(*t*), from the baseline activity **r**_0_. We take *r* = *f*(*x*) = *x* for the mathematical analysis, and compare to networks with richer dynamics using a known non-linear function^4,8^. In the linear case, the network dynamics can be described using the eigenvalues, *λ_i_*, and eigenvectors, **v***_i_*, of the weight matrix **W** (with *i* = 1, −, *N*; *N* the number of neurons in the network). To quantify whether and by how much the network can amplify specific inputs, we calculate the norm of the rate vector, ‖**r**(*t*)‖, by decomposing it in the directions of the eigenvectors of **W**,

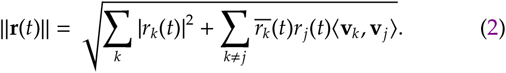

Here, *r_k_*(*t*) = *α_k_e*^(*λ_k_*–1)*t*^ is the solution of the system along the direction of the eigenvector **v**_*k*_, which is associated with the eigenvalue *λ_k_* (*α_k_* is a constant, uniquely determined by the initial condition). In a stable regime, Re(*λ_k_*) < 1, ∀*k*, the system exhibits a single fixed point that represents the baseline activity. An increase of the response norm, ‖**r**(*t*)‖, with respect to the norm of the initial condition, ‖**r**(*t*_0_)| (here always normalised to 1), defines the phenomenon of transient amplification. A necessary condition for this to happen is the non-normality of **W**, 〈**v***_k_*, **v***_j_*〉 ≠ 0, for some *j*, *k*, i.e., the eigenvectors do not form an orthogonal basis^11^. To explore regimes of transient amplification, we thus focus on matrices of the form 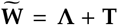 (Fig. 1B), with the diagonal, **Λ**, containing the eigenvalues^10–12^, and the strictly upper triangular part, **T**, representing the feedforward structure between patterns of activation. Note that **Λ** contains 2 × 2-blocks around the diagonal to accommodate for complex eigenvalues in real-valued matrices. The real parts of the eigenvalues are on the diagonal and the imaginary parts lie on the off-diagonal entries of the 2 × 2 blocks^19^. We create **Λ** by sampling the real and imaginary parts of the eigenvalues from different distributions, but keeping the number of complex versus real eigenvalues constant (here 3% real). The imaginary distribution needs to be symmetric with respect to zero (a condition imposed by the conjugacy of the complex eigenvalues), while the real distribution must be below 1 (and is here always set to have 0.5 as a supremum) for stability reasons. We create **T** in two different ways: from the Schur decomposition of a stability optimized circuit^20^, or sampled from a uniform distribution. We scale the norm of **T** after its structure is fixed.

We start our investigation of how the eigenspectrum affects the dynamics by drawing both real and imaginary parts from uniform distributions with diameters *d*_im_ and *d*_re_, respectively (Fig. 1C–F, top left). To quantify the dynamical response of the network, we find an optimal orthogonal basis of initial conditions, 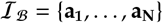, ordered according to their evoked energy, 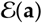^21^. To make sure that the evoked energy is due to an amplified response rather than merely a slower exponential decay, we compute the maximum value of the norm of the firing rate vector, for all vectors in 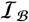 (Fig. 1C–F, bottom left).

With broad distributions, the system can slightly amplify a few conditions (Fig. 1C). When the range of the real–part distribution is decreased and pushed towards 0.5, the resulting network produces stronger amplification (Fig. 1D). This can mainly be attributed to the fact that the eigenvalues have now larger real parts and hence longer decay envelopes. Indeed, clustering away from 0.5 leads to less amplification (not shown).

More surprisingly shrinking, instead, the imaginary distribution also leads to more amplification (Fig. 1E), and shrinking both produces very large amplification that in the non-linear case lasts for a long time, approximating timescales of working memory dynamics (Fig. 1F). Additionally, the percentage of conditions that are amplified is considerably increased, i.e., the ability of such a network to amplify orthogonal initial conditions is enhanced. Note that splitting and clustering the (positive and negative) imaginary parts away from zero gives rise to slightly different amplification regimes that also depend on the linearity of the system^19^.

When we study the effects of the imaginary and real distributions more systematically, we find that the shape of the real distributions^19^ has minimum effect on the amplification (Figs. 2Ai and 2Aii). Amplification emerges from the nonnormality of **W**, which can be partly quantified by the angles between the eigenvectors (Eq. 2)^19^; if more pairs have overlaps, the matrix will be more non-normal. The imaginary distribution changes the geometry of the eigenvectors (Fig. 2Aiii), providing a mechanism for its drastic effect on the amplification in these networks (Figs. 2Ai and 2Aii). This is a surprising effect given that we do not alter the feedforward norm nor the decay envelopes at all.

**FIG. 2.**
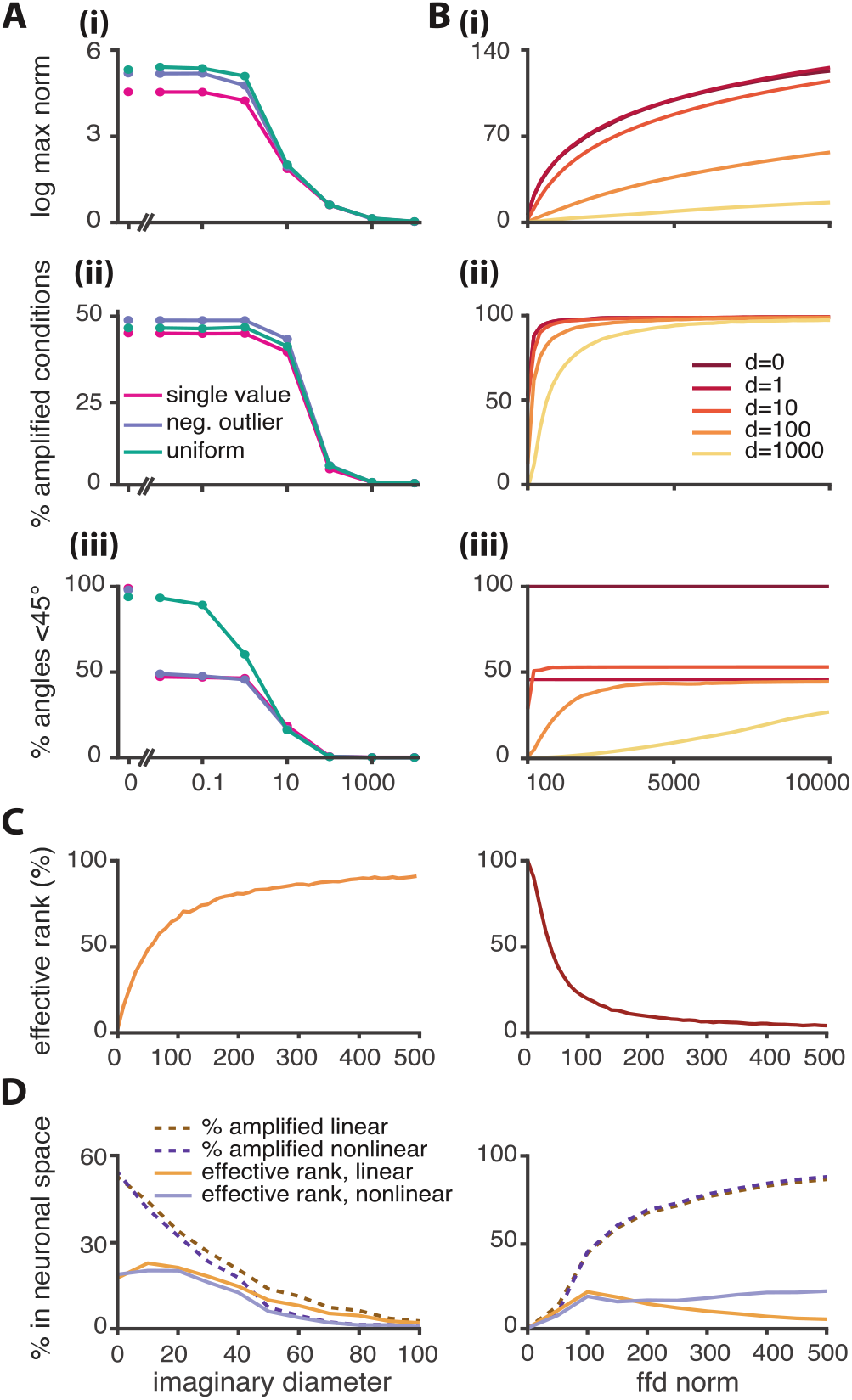
Effects of manipulating the spectrum and feedforward norm. **A**, **(i)** Maximum response norm for the preferred initial condition, **(ii)** Percentage of directions whose norm is amplified more than 50% and **(iii)** The percentage of angles (between pairs of eigenvectors) that are less than 45°. Every line is a function of the imaginary diameter. We plot three real distributions. Pink; a single valued real distribution in which all real parts are equal to zero. Purple; a distribution with a large real negative outlier at −20 and the rest of the real eigenvalues distributed between the values 0 and 0.5. Green; a uniform distribution in which all real parts are distributed uniformly in the interval (−0.5,0.5)^19^. **B**, **(i-iii)** Same as A, but plotted as a function of the feedforward Frobenius norm. Different colours correspond to 5 different spectra; all spectra have fixed single-valued real distributions (equal to zero) and different imaginary diameters. **C**, The effective rank of the eigenvector matrix **V** of 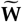 as a function of the imaginary diameter (orange) or the ffd norm (red). **D**, Amplified directions and effective rank of the matrix 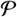 (see text) in the linear and nonlinear cases. The feedforward structure is random, from a uniform distribution, and the real distribution is uniform on (−0.5,0.5). **(i)** As a function of the imaginary diameter with feedforward norm equal to 75. **(ii)** As a function of the feedforward Frobenius norm, with imaginary diameter equal to 20.

The feedforward norm is more directly linked to the non-normality^11^, and as expected, it increases both the norm of the maximum response (Fig. 2Bi), and the percentage of amplified conditions (Fig. 2Bii), for larger values. The percentage of eigenvector pairs with small angles also grows with increasing feedforward norm strength (Fig. 2Biii). Interestingly, there is a saturating point that depends on the imaginary distribution. Once the number of pairs saturates, increased amplification is mainly due to the increased matrix norm, ‖**W**‖, indicating that the eigenvector pairwise angles are not sufficient to explain the behaviour of the network. Next we study the relative position of the directions of the overlaps in state–space.

If most eigenvectors are pointing in similar directions, the dynamics will be biased towards these directions too. This does not mean that **W** or the eigenvector matrix **V** are not full rank–on the contrary, they almost always are. What it means is that, in order to quantify the global eigenvector geometry, we have to use the effective rank of **V**. The effective rank of **V** measures the average number of significant dimensions in its range, and is formally defined as the exponential of the spectral entropy of its normalised singular values^22^. Specifically, if *σ*_1_, *σ*_2_, ⋯, *σ_N_* are the singular values of **V**, and 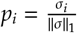, with 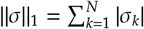, then:

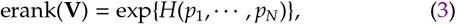

where *H*(*p*_1_, ⋯, *p_N_*) is the Shannon entropy, i.e., 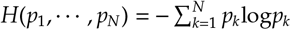. The effective rank of **V** is indeed small in the highly amplifying regimes (Fig. 2C), revealing an underlying duality between amplification and output dimensionality.

The consequence for the dynamics is that, even though the system may amplify many initial conditions, they nevertheless evolve in the same low dimensional subspace. To identify the dimensionality of this subspace we compute the effective rank of the matrix 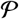 which is constructed as follows: the *j*-th column of 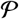 is the first principal vector of the dynamics, given the *j*-th amplified initial condition of the 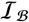 basis^19^. We find that there is a discrepancy between the number of amplified directions and effective rank when the system produces large amplifications (Fig. 2D). This suggests that the dynamical responses evoked by orthogonal initial conditions evolve in the same subspace. This also means noise will be amplified in the same direction as the signal. Additionally, different initialisations would potentially lead to similar linear readouts. There is thus a trade-off between the number of amplified conditions, i.e., the capacity, and the noise robustness of the system.

To summarise, the system can be described by three regimes of amplification: weak; short transient; and long transient. In the weak case, the eigenvectors are effectively orthogonal to each other but span the entire output space equally. In the short transient regime, there is a good balance between amplification of orthogonal inputs and diversity in the responses. In the long transient, many initial conditions are amplified but the responses lie in the same low-dimensional subspace. Moreover, we found that the mechanism behind the different regimes of amplification depends on the difference between the norm of the eigenspectrum and the norm of feedforward structure (Fig. 3A). Indeed, when we fix the norm of **W**, and distribute a–continuously decreasing– percentage of this norm on the diagonal and the rest on the feedforward structure, the network transitions from weakly to strongly amplifying (Fig. 3B). Thus, it’s the relation between the diagonal (representing the spectrum) and feedforward parts of the matrix that shapes the dynamics of the network.

**FIG. 3.**
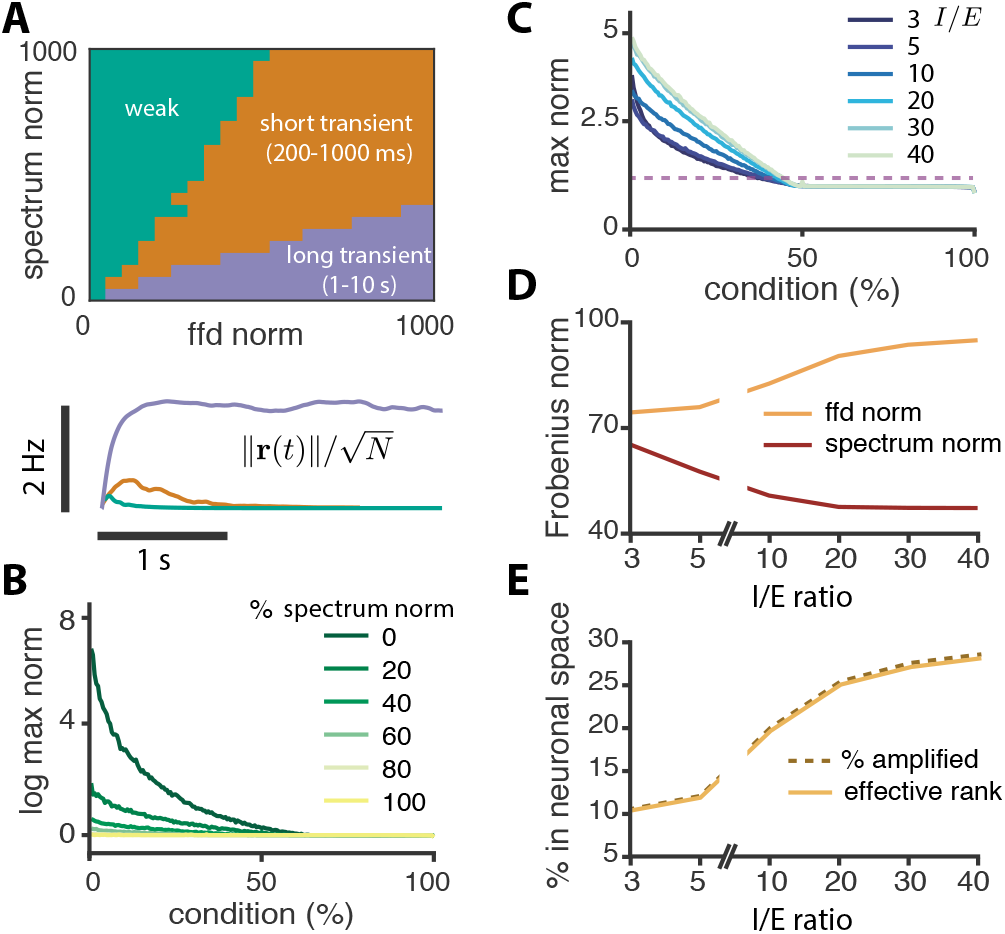
Amplification regimes and the effect of inhibitory dominance in Dalean matrices. **A**, Timescale of the response (time for which ‖**r**(*t*)‖ ≥ 1) in the nonlinear network, parametrised by the norms of the spectrum and feeforward structure. Bottom: three representative examples from each regime. **B**, Maximum norm of the dynamical response per initial condition for different percentages of the norm assigned to the spectrum, ranging from a matrix whose entire norm is assigned to the spectrum (yellow; 100% case, normal matrix) to a matrix whose entire norm is assigned to the feedforward part (dark green; 0% case, nilpotent matrix). **C–E**, Simulations for a connectivity matrix satisfying Dale’s law (50% excitatory and 50% inhibitory neurons). An initial spectrum of radius 10 with random connections but global inhibitory dominance of strength *I/E*. Following the optimisation algorithm used in SOCs^8^, we stabilise the network keeping the initial *I/E* ratio. **C**, The amplification landscape for different *I/E* ratios. Purple dotted line corresponds to a response norm that is 50% larger than the norm of the initial condition. **D**, The evolution of the spectrum and feedforward norms for different values of *I*/*E* in the corresponding real Schur transformation. **E**, Percentage of amplified conditions and effective rank of the corresponding matrix 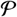 (defined in text) in the linear case.

Next, we can investigate neuronal networks with certain biological constraints. First, we consider the effect of nonself loops in the connectivity matrix. Neurons are not typically structurally connected to themselves, which means that the trace of the weight matrix of such a network is equal to zero. This unfolds as follows: 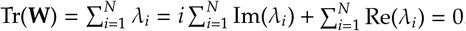. Given that 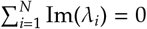 due the conjugacy of the eigenvalues, the weight matrix **W^B^** of an upper triangular matrix without self-loops requires 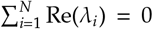. This, together with the stability constraint, max(*α*(*W*)) = 0.5, bounds the real distribution from below and above, restricting it to a very limited range. The observation explains why the spectrum of the SOC has the pancake shape after optimisation, i.e., not only the positive eigenvalues but also the negative are pushed towards the stability line after optimisation. This result also highlights the importance of the imaginary spectral manipulations: if the real distribution is limited, the imaginary spectrum must probably carry important information^5^, and its role is likely critical for shaping the dynamics.

As a last application, we explore how to navigate the regimes of transient amplification in networks with excitatory and inhibitory neurons, i.e., satisfying Dale’s law. In this case, we have to design a biological matrix **W**^B^ which has a real Schur transformation with *large* feedforward and *small* diagonal norm. We find that a mechanism to control this in the short transient regime, is the strength of global inhibition. Larger inhibitory global strength leads to more amplified conditions and also to larger amplification per condition (Fig. 3C). By assigning larger values to the inhibitory weights, the feedforward norm increases and the spectrum norm decreases (Fig. 3D). Finally, the new amplified conditions induced by the strongest inhibition do not share their first principal component directions in their dynamical responses, i.e., the noise robustness of the system is not compromised in this case (Fig. 3E). This is possible because we are still in the short transient regime; the long transient regime cannot be reached by solely increasing the global inhibitory strength^19^.

In this letter we used upper triangular matrices as abstract representations of the dynamical properties of a connectivity matrix, to control the quantities that are relevant for the neural dynamics in the transient amplification regime. Although transient non-normal amplification has been previously studied^4,6,8,9,11^, the entire dynamical regime that can be spanned by this kind of networks had not been explored. Usually, any alteration in the weights of a neuronal connectivity matrix has obscure effects on the spectrum and non-normality. By by-passing, temporarily, the connectivity matrix and focusing on a hypothetical Schur transformation, we found new dynamical regimes of large amplification that had not been reported before. We also showed that the amount of transient amplification a network can produce can be controlled by the ratio between the norms of the spectrum and hidden feedforward structure that the Schur transformation unveils. Moreover there is a trade–off between the capacity and noise robustness of those systems. The source of amplification, i.e., the overlaps of the eigenvectors, inevitably restrict the subspace in which the dynamical outputs evolve. Finally, we found that stronger global inhibitory dominance helps navigate amplification regimes in networks that satisfy Dale’s law. Our work opens the door for the exploration of new questions related to neuronal dynamics, such as how the structure – besides the norm – of the feedforward part as well as how non-uniform imaginary distributions affect the dynamics.

We thank F. Zenke for his comments, and especially his contribution on the effect of the self-loops on the spectrum. We also thank G. Hennequin, Ken Miller, and W.F. Podlaski for helpful comments on the work and manuscript. This work was supported by a Wellcome Trust and Royal Society Henry Dale Research Fellowship (WT100000, TV), a Wellcome Senior Research Fellowship (214316/Z/18/Z; GC, EJA and TPV), and a Research Project Grant by the Leverhulme Trust (RPG-2016-446; EJA).

## Supplementary material

### Details for upper triangular matrix setup

To construct the upper triangular matrices we start with an example of a connectivity matrix known to create strong non-normal amplification, i.e., the Stability Optimised Circuit (SOC)^2^. We take the real Schur transformation of this connectivity matrix. In this form, the matrix is upper triangular with some 2 × 2 blocks on the diagonal. These blocks have real entries and their eigenvalues are the complex eigenvalues of the initial matrix (a pair of conjugates). We fix the triangular part that is not involved in the eigenvalue blocks. In most of the manipulations of the feedforward coupling, we only change the norm of this feedforward part of the matrix by scaling all its entries accordingly. In some cases (for control) we also chose the feedforward connections to be drawn from a uniform distribution on the interval (−1,1), and scale the norm accordingly as well. For the manipulation of the spectrum we construct our distributions by hand. Specifically, if we want the matrix to have the pair of complex eigenvalues *α* ± *ωi* in its spectrum, we add the block

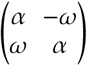

along the diagonal. For real eigenvalues, we just add the corresponding real value on the diagonal.

### Why upper triangular?

The idea behind the use of an upper triangular matrix arises from the real Schur decomposition. Given a connectivity matrix **W**, one can find the eigenspectrum using the basis of eigenvectors. However, the non-normality of the matrix is lost under this linear transformation. Since we are especially interested in the dynamical regime of transient amplification we have to go beyond the spectrum, and a better way to access the nonnormality is to use its Schur decomposition. Indeed, any square matrix is unitarily equivalent to an upper triangular one, and by definition, the minimum over all such decompositions norm of the strictly upper triangular part is its non-normality index. We follow the same idea, but use instead the real Schur transformation. The advantage is that we still have in our hands a real-valued matrix. The disadvantage is that we now have to deal with 2 × 2 blocks along the diagonal. However, the important thing is that **W** is still orthogonally equivalent to its real Schur transform. This means that the non-normality quantity we are interested in is still preserved, i.e., the dynamical characteristics of transient amplification between the two matrices are not qualitatively different.

### Details on the real distributions

The real distributions we compare in the main text are the following:

- **A single valued distribution**: all real parts are the same and equal to a fixed value. In the simulations shown in the main text, this value is taken to be zero for comparison reasons, and also to agree with the (subsequently introduced) idea of the zero trace condition. In figure S1A we compare to cases where this fixed value is equal to −0.5 and 0.5.
- **A distribution with a negative outlier**: in this construction, we add a negative outlier at a specific point. Because we require that the sum of all real parts is equal to zero, we have to set some real parts to be equal to 0.5. The number of real parts set at 0.5 depends on the value of the outlier. The rest of the real values are set to zero.
- **A uniform distribution on the interval** (−0.5,0.5): all real parts, except for the last one, are distributed uniformly between the values −0.5 and 0.5. As before, because of the zero trace condition, we have to add a small outlier (the last real eigenvalue) to complement for the non-zero sum of the rest of the values.

In all cases the pairings with the corresponding imaginary parts are random – except for forcing the conjugacy of eigenvalues, that is, we make sure that the same real part is paired with conjugate imaginary parts. All simulations are run for 200 realisations, with respect to the randomness of the imaginary distribution, and final quantities are averaged across all realisations for plotting.

### Details on the feedforward structure

In some simulations, the feedforward structure of the upper triangular matrices was taken to be equal to the upper triangular part of the Schur transform of a fixed stability optimised circuit. In other simulations, the upper triangular part of the matrix was simply drawn from a uniform distribution. In Figure S1B we compare the results for both structures when the real and imaginary distributions are identical.

### Eigenvector overlaps

Recall that the eigenvectros are, in general, complex, in conjugate pairs and that in order to compute the overlap of the eigenvectors we need to consider their inner product. The inner product of two complex numbers is defined as

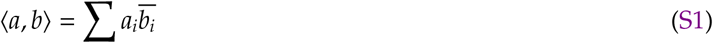

and the angle between two complex vectors is given by

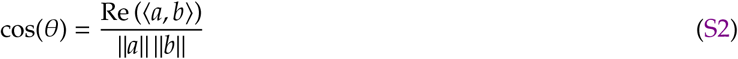

Therefore, to compute the angles between the eigenvectors we use Equation S2. In particular, we normalise the eigenvectors to unit norm and compute all pairwise angles. Finally, since cos(*π* – *θ*) = −cos(*θ*), when computing the percentage of small eigenvector overlaps (i.e. less than 45°), we consider as angle the minimum angle between *θ* and *π* – *θ*. We would like to note here that non-normality depends on the complex inner product between eigenvectors, and not only its real part. However, we have chosen to compute this more intuitive version of an angle between two complex vectors (which is commonly used in the literature) as a characterisation of the amplification dynamics.

### Imaginary clustering at different points

To understand whether the surprising effect of the imaginary spectrum is due to the clustering of the eigenvalues we checked what happens when the imaginary clusters are not around zero, but at e.g. the points {100, −100} (Fig. S2A inset). In this case the linear responses exhibit an interesting phenomenon, resembling the beats in acoustics (Fig. S2A). Because the frequencies are close to each other (due to the clustering), the amplitudes of the different neuronal responses are superimposed when phased, to create a response of very high amplitude (which by our definition would count as amplification). Moreover, the differences in the frequencies create an envelope that is modulating this amplitude over time. On the other hand, the nonlinear responses fail to capture most of the interesting dynamics seen linearly and do not amplify to the same extent (Fig. S2B). The very high frequency makes it impossible for any potentially amplifying mode to drive the rest of the nodes and create a big amplified response. Because of this discrepancy between linear and nonlinear behaviour, we will not consider these regimes as amplifying for the purposes of this letter. It is worth noting that similar behaviour to the ±100 example is seen when clustering the imaginary spectrum at different nonzero values (Fig. S2C).

### Dimensionality of dynamics–effective rank of eigenvector matrix

Here we briefly explain the intuition behind the effective rank of the eigenvector matrix **V**. This is understood as the number of significant dimensions in the range of a matrix. For example, if the effective rank is equal to *κ*, then a random trajectory in the range of **V** is sufficiently approximated by *κ* dimensions. The fact that the effective rank of the eigenvector matrix is small indicates that there are a few prevalent directions in the space spanned by the eigenvectors, which indicates that dynamical trajectories will be biased towards a small subspace of the entire eigenvector space. This is further explored and verified with the computation of the dynamical matrix 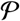 defined below.

### Construction of the matrix 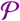

We constructed the matrix 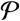 to understand how correlated the dynamics of the network are given different initial conditions. This matrix represents the prevalent directions of the dynamics, given different initialisations. This is done as follows: after having identified the optimal orthogonal basis of initial conditions 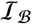, we initialise the network at each of the vectors in this basis, one at a time. For each such vector, if the induced dynamics are amplified, i.e., if the norm of the rate vector is at some point in time larger than 1.5 (the initialisation vectors have always unit norm), then we perform Pricipal Component Analysis on the dynamics. More specifically, we compute the eigenvectors of the covariance matrix of the neuronal dynamics for each of these simulations. From these eigenvectors we only consider the eigenvector corresponding to the largest eigenvalue and store it as a column in the matrix 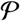. Once we have initialised the network at all vectors in 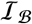 we are left with a *N* × *M* matrix 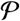. The number *M* is the same as the number of conditions that lead to an amplified response and provides a maximum bound for the effective rank of matrix 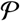.

The effective rank of 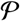 thus gives us the effective dimensionality of the space spanned by the columns of 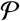. If the effective rank is less than the the number of columns, we can deduce that orthogonal initial conditions have first principal components that are close to each other in state-space. This implies that the initial network amplifies orthogonal initial conditions along the same low dimensional subspace.

### Inhibitory dominance

To inutitively understand why the strength of inhibitory dominance might facilitate amplification regimes, we can first model global inhibitory dominance abstractly in a matrix 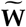 that is upper triangular. We do this by adding a real negative outlier to the real spectral distribution. Interestingly, the existence of the negative outlier together with the zero trace condition has a very interesting effect. The larger the value of the outlier (in absolute value), the bigger the amplification (Fig. S3A) and the number of amplified directions (Fig. S3B). On one hand, this can be explained by the fact that more real parts are pushed to the right, creating longer decay envelopes, hence prolonging the time for the hidden feedforward structure to be amplified. However this is not the sole source of the increased amplification; a larger negative outlier has an additional non-intuitive effect on the geometry of the eigenvectors, i.e., it gives rise to larger eigenvector overlaps (Fig. S3C).

We confirm the above result in the main text, with a biologically plausible matrix, **W**^B^ satisfying Dale’s law – columns have either only positive or negative entries (Fig. 3C–E). Taking into account the effect of the negative outlier, we hypothesized that assigning the matrix strength in the inhibitory weights will result in this *good* real Schur decomposition. Larger inhibitory weights can lead to a larger global inhibitory dominance ratio, hence to a larger negative outlier and therefore to more amplification according to the theoretical, upper triangular matrix results (Fig. S3). To highlight this finding, we note that in the example shown in Fig. 3C–E, when the inhibitory to excitatory ratio is large, *I/E* = 40, the strength of every nonzero excitatory-to-excitatory connection is 0.08, and yet the network is capable of stronger amplification compared to when *I/E* = 3 in which the nonzero excitatory-to-excitatory weights are set to 1.05. This also explains why we can’t reach the long transient regime by increasing the inhibitory strength. Since the overall norm of the matrix stays the same (for comparison reasons), increasing more than that the inhibitory dominance would necessarily decrease even further the excitatory weights. Therefore, the amplification power of the network through this mechanism eventually saturates before reaching the long transient regime.

## Supplementary figures

**FIG. S1.**
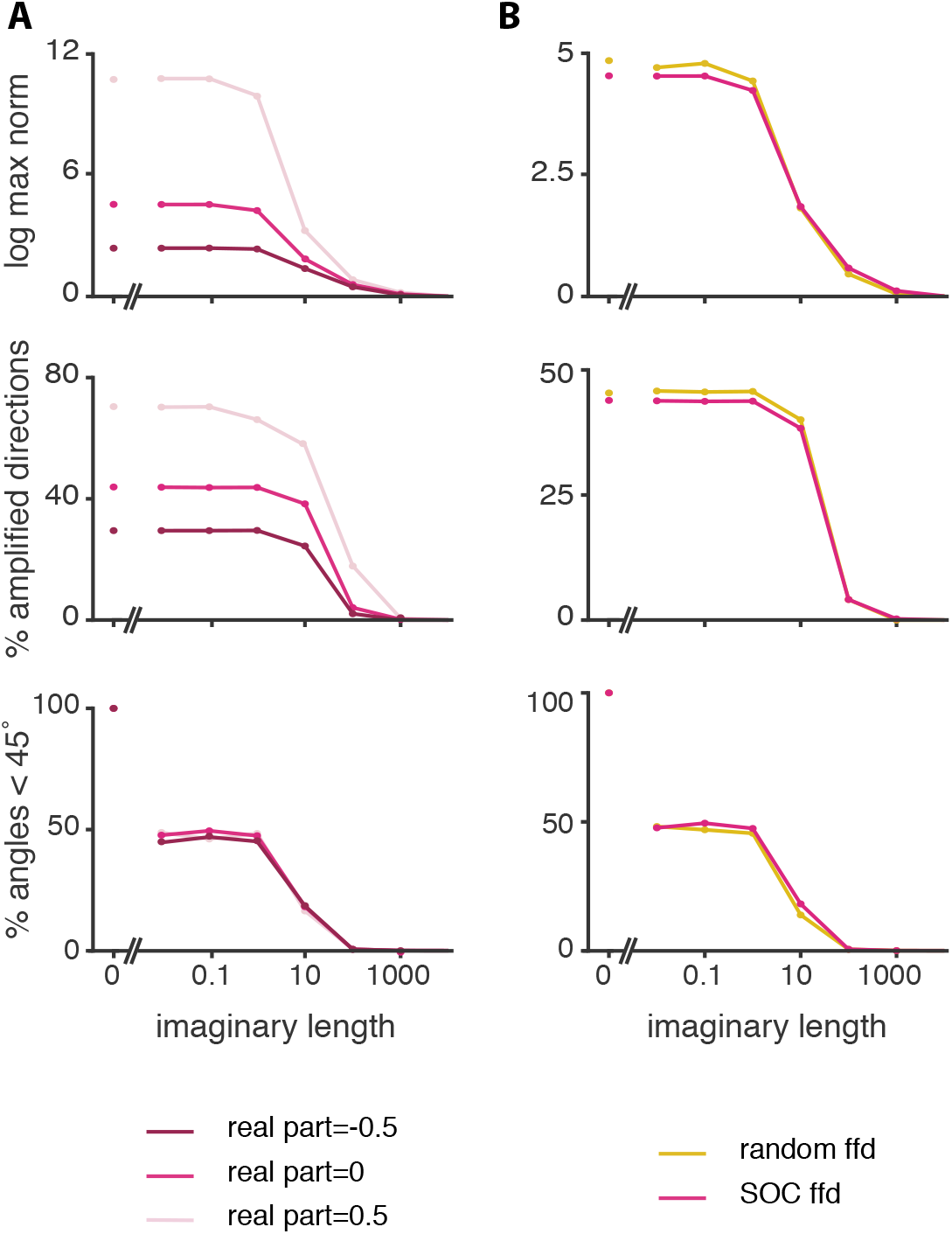
Variations on the real and ffd distributions. **A**, Exploring the single valued real distribution. We compare results for three real values: −0.5,0 and 0.5. Top: maximum response norm for preferred initial condition. Naturally a larger real part leads to more amplification as the decay envelope becomes slower. Middle: % of amplified conditions that are amplified by at least 50%; this is also affected by the value of the real part indicating that the the amplification landscape changes its shape in a uniform way. Bottom: the percentage of pairwise eigenvector angles is independent of the real value, i.e., the increased amount of amplification is mainly a result of the slower decay times. Results in all cases are qualitatively similar in their evolution with respect to the imaginary radius. **B**, Comparing results for two different feed-forward structures. One is the feed-forward structure taken from the corresponding ffd part of a matrix constructed using the SOC algorithm (pink). The other has a uniform ffd entry distribution, with overall ffd norm equal to the SOC one (yellow). In both cases the spectra are identical and correspond to the spectral distribution of the pink curve from A, i.e., single real value at zero, varying imaginary range represented on the x-axis.

**FIG. S2.**
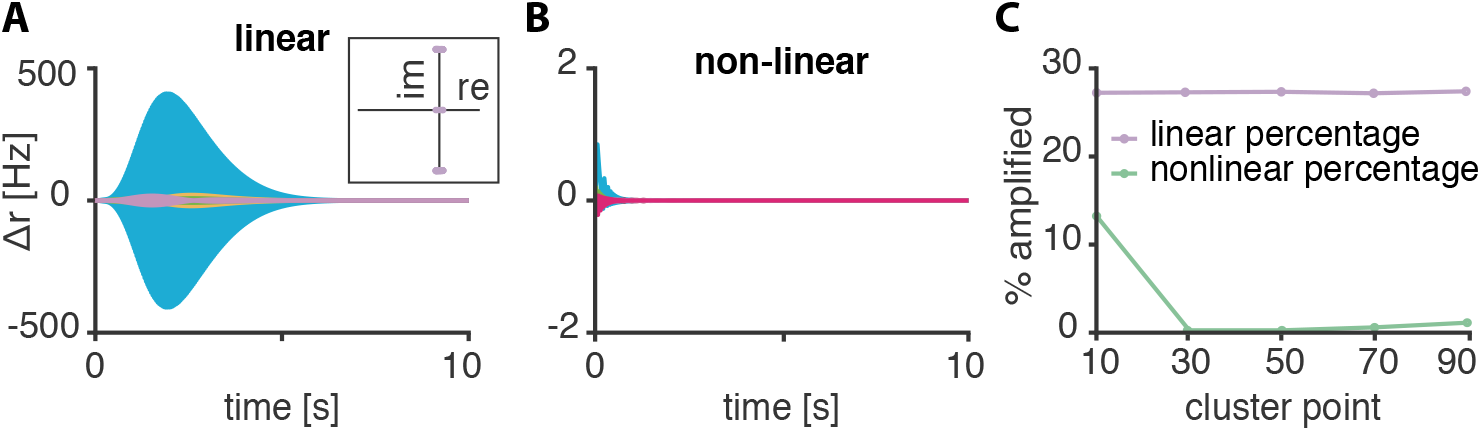
Imaginary clustering at different points. Dynamical responses of a spectrum that is clustered around the points 100 and −100 with respect to the imaginary axis. The imaginary radius around these points is 0.5. The real distribution is uniform on the interval (−0.5,0.5). **A**, Linear dynamical response; the network shows an amplified response, effectively due to superposition of almost identical frequencies. **B** Nonlinear responses are not amplifying; the very high frequency together with the saturation point prevents the network’s modes from driving each other in order to create an amplifying response. **C** Clustering at other finite points, {±10, ±30, ±50, ±70, ±90} shows the same discrepancy between the linear and nonlinear behaviour; as measured by the percentage of amplified responses.

**FIG. S3.**
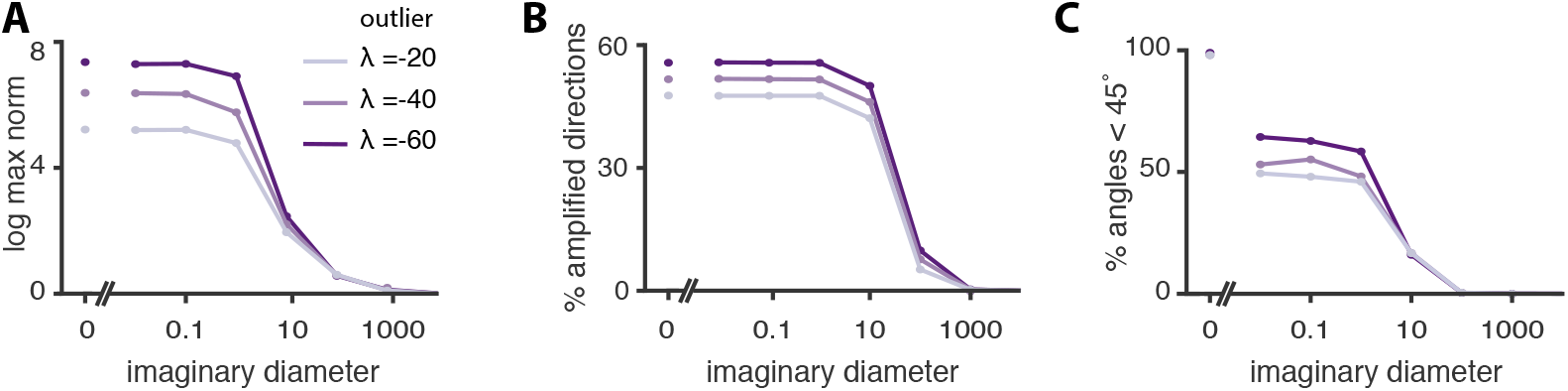
Global inhibition in the upper triangular setting. **A**, Maximum response norm for the preferred initial condition, **B** Percentage of directions whose norm is amplified more than 50% and **C** The percentage of angles, between pairs of eigenvectors, that are less than 45°. Every line is a function of the imaginary diameter which ranges from *d* = 0 to *d* = 10000. Different colours correspond to three different real distributions in the following way: All distributions have a large negative outlier (hypothetically arising from some sort of globally dominating inhibition) and satisfy the no self-loop condition; the position of the negative outlier determines the colour. Light purple; *λ* = −20. Medium purple; *λ* = −40. Dark purple; *λ* = −60. Larger in amplitude outlier creates more amplification.

## References

[1] T. P. Vogels, K. Rajan, and L. F. Abbott, Neural network dynamics, Annu. Rev. Neurosci. 28, 357 (2005).

[2] K. Rajan and L. F. Abbott, Eigenvalue spectra of random matrices for neural networks, Physical Review Letters 97, 188104 (2006).

[3] D. Sussillo and L. F. Abbott, Generating coherent patterns of activity from chaotic neural networks, Neuron 63, 544 (2009).

[4] J. P. Stroud, M. A. Porter, G. Hennequin, and T. P. Vogels, Motor primitives in space and time via targeted gain modulation in cortical networks, Nature Neuroscience 21, 1774 (2018).

[5] L. Susman, N. Brenner, and O. Barak, Stable memory with unstable synapses, Nature Communications 10, 4441 (2019).

[6] G. Bondanelli and S. Ostojic, Coding with transient trajectories in recurrent neural networks, PLOS Computational Biology 16, e1007655 (2020).

[7] V. Mante, D. Sussillo, K. V. Shenoy, and W. T. Newsome, Context-dependent computation by recurrent dynamics in prefrontal cortex, Nature 503, 78 (2013).

[8] G. Hennequin, T. P. Vogels, and W. Gerstner, Optimal control of transient dynamics in balanced networks supports generation of complex movements, Neuron 82, 1394 (2014).

[9] E. D. Remington, D. Narain, E. A. Hosseini, and M. Jazayeri, Flexible sensorimotor computations through rapid reconfiguration of cortical dynamics, Neuron 98, 1005 (2018).

[10] G. Hennequin, T. P. Vogels, and W. Gerstner, Non-normal amplification in random balanced neuronal networks, Physical Review E 86, 011909 (2012).

[11] B. K. Murphy and K. D. Miller, Balanced amplification: a new mechanism of selective amplification of neural activity patterns, Neuron 61, 635 (2009).

[12] M. S. Goldman, Memory without feedback in a neural network, Neuron 61, 621 (2009).

[13] M. M. Churchland, J. P. Cunningham, M. T. Kaufman, S. I. Ryu, and K. V. Shenoy, Cortical preparatory activity: representation of movement or first cog in a dynamical machine?, Neuron 68, 387 (2010).

[14] M. M. Churchland, J. P. Cunningham, M. T. Kaufman, J. D. Foster, P. Nuyujukian, S. I. Ryu, and K. V. Shenoy, Neural population dynamics during reaching, Nature 487, 51 (2012).

[15] K. C. Ames, S. I. Ryu, and K. V. Shenoy, Neural dynamics of reaching following incorrect or absent motor preparation, Neuron 81, 438 (2014).

[16] M. Shafi, Y. Zhou, J. Quintana, C. Chow, J. Fuster, and M. Bodner, Variability in neuronal activity in primate cortex during working memory tasks, Neuroscience 146, 1082 (2007).

[17] O. Barak, M. Tsodyks, and R. Romo, Neuronal population coding of parametric working memory, Journal of Neuroscience 30, 9424 (2010).

[18] C. R. Hussar and T. Pasternak, Memory-guided sensory comparisons in the prefrontal cortex: contribution of putative pyramidal cells and interneurons, Journal of Neuroscience 32, 2747 (2012).

[19] Supplemental Material with: (i) Details for upper triangular matrix setup; (ii) Why upper triangular?; (iii) Details on the real distributions; (iv) Details on the feedforward stricture; (v) Eigenvector overlaps; (vi) Imaginary clustering at different points; (vii) Dimensionality of dynamics–effective rank of eigenvector matrix; (viii) Construction of the matrix P; and (ix) Inhibitory dominance.

[20] Without loss of generality, the fixed structure is identical to the strictly upper triangular part of a Stability Optimised Circuit (SOC)^8^; a network that is known to amplify and whose amplification statistics (degree of amplification and number of amplified conditions) can be used as a benchmark. Using a random uniform distribution in T with the same feedforward norm does not change the results qualitatively^19^.

[21] If **x**(*t* = 0) = **a**, ‖**a**‖ = 1, then 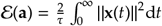, from Hennequin et al.^8^.

[22] O. Roy and M. Vetterli, The effective rank: A measure of effective dimensionality, in 2007 15th European Signal Processing Conference (IEEE, 2007) pp. 606–610.

